# *stPipe:* A flexible and streamlined R/Bioconductor pipeline for preprocessing sequencing-based spatial transcriptomics data

**DOI:** 10.1101/2025.04.16.649254

**Authors:** Yang Xu, Callum J. Sargeant, Yue You, Yupei You, Shian Su, Changqing Wang, Luyi Tian, Yunshun Chen, Matthew E. Ritchie

## Abstract

Spatial transcriptomics technology has developed rapidly in recent years, with various sequencing-based platforms such as 10x Visium, Slide-seq and Stereo-seq becoming widely used by researchers. Each platform brings its own set of protocols and customised data analysis pipelines which presents challenges when the goal is to obtain uniformly preprocessed data that is conveniently formatted for downstream analysis. To address the need for simpler, open-source solutions that deal with sequencing-based spatial transcriptomics (sST) data from different platforms, we present `stPipe`, a comprehensive and modular pipeline for analysing sST data. `stPipe` handles various analysis steps including (i) data processing from raw paired end FASTQ files to create a spatially resolved gene count matrix; (ii) the collation of relevant quality control metrics during preprocessing to ensure unwanted artefacts can be filtered from further analysis; and (iii) the adoption of standardised data storage containers to allow results to be easily passed on to a wide range of downstream analysis packages tailored to different goals (such as clustering, cell-cell communication analysis and differential expression analysis). `stPipe` is implemented as an R/Bioconductor package that builds upon functionality in the `scPipe` software, and offers a flexible preprocessing pipeline that can manage data from all current main-stream sST plaforms. A key use case for `stPipe` is in methods benchmarking, and we demonstrate how the uniform processing of sST data collected on reference tissue samples from the *cadasSTre* and *SpatialBenchVisium* projects is made easier, allowing comparisons between different technology platforms and downstream analysis tools. Our framework thus aims to advance the standardization and optimization of spatial transcriptomics analyses, fostering collaboration and innovation within the research community.

## Introduction

The rapid development of spatially resolved transcriptomics technology over the past 10 years has enabled a more comprehensive understanding of the patterns of gene expression and cellular communication across tissues (1). Spatial transcriptomics (ST) has seen broad application in investigating tissue development, spatially resolved biomarker identification, cell state and fate determination across tissues in a wide range of research fields, including immunology, developmental biology, oncology and neuroscience (2). Various ST platforms that employ distinct methodologies have been developed. These can be broadly classified into two main categories: imaging-based spatial transcriptomics (iST) and sequencing-based spatial transcriptomics (sST) which differ fundamentally in their strategies for determining spatial localization and quantifying mRNA abundance (3). Commercially available iST technologies such as Xenium (10x Genomics) and MERSCOPE (Vizgen) leverage fluorescent signatures and their intensity to measure target genes and their abundance via highly multiplexed single molecule fluorescence in situ hybridization (smFISH). While imaging-based methods excel in spatial resolution, they are typically limited by the number of transcripts that can be simultaneously profiled (4). In contrast, commercial sST platforms like Visium (10x Genomics) and Stereo-seq (BGI) depend on spatial barcodes arranged either deterministically or randomly on an array to reconstruct the spatial position of the target gene within the tissue. These sST methods then apply next-generation sequencing (NGS) to the spatial libraries constructed to ascertain their expression level. Sequencing-based methods tend to sacrifice spatial resolution, but allow genome-wide gene quantification.

10x Genomics Visium is one of the most popular and widely used ST platforms over the past 5 years (3). 10x Genomics provides two Visium capture chemistries which enable either fresh frozen (FF) tissue samples to be profiled using polyA-based capture of RNA or the analysis of Formalin-fixed paraffin-embedded (FFPE) samples based on RNA templated ligation (RTL) of pairs of gene target probes. Both of these approaches use slides covered with 55µm diameter spots with a center-to-center distance of 100µm between horizontal spots (5). For FFPE samples, pre-defined human (or mouse) transcriptome-centric probe-sets and spatial barcode allow lists enable Visium to capture both gene expression and spatial localization information simultaneously. For polyA-based FF samples, splicing-aware alignment of transcripts is performed for reads to map them to the reference genome. To accurately align ST sequencing data with the tissue image and perform pixel computation for each spot, fiducial alignment and tissue detection are required. To simplify the original Visium workflow, 10x Genomics developed the Visium CytAssist instrument to allow transfer of the tissue section from a standard glass slide onto the Visium slides, allowing the use of either pre-sectioned tissue or FFPE blocks (2). The recently released Visium HD platform enables single cell-scale resolution via the tiled arrangement of millions of 2 × 2µm barcoded squares that are typically binned (into 8 × 8µm or 16 × 16µm regions) for further analysis. This represents a greater than 4-fold increase in spatial resolution over regular Visium technology. A practical example of where this enhanced resolution improves the biological insights gained was recently demonstrated in the study of normal colon mucosa, with Visium HD able to capture the characteristic hole pattern via the binning of barcoded squares while the lower resolution Visium data could not recover this expected detail (6).

BGI’s DNA Nanoball (DNB) technology is used by Stereoseq (SpaTial Enhanced REsolution Omics-sequencing) to recover data on the STOmics-GeneExpression Chip at up to nanometer level resolution (7). DNB is generated by rolling circle amplification of circularized DNA oligonucleotides, forming spherical structures. Each DNB carries a 25-nt co-ordinate identity (CID) used to identify its spatial position during RNA capture and sequencing. The spatial location of each DNB is recorded with a diameter of 0.2µm and a center-to-center distance of 0.5µm to generate sub-cellular single-cell resolution data. Compared to 10x Visium technology, Stereo-seq can be analysed with less reliance on scRNA-seq data to perform deconvolution of the cell type mixtures that are present in data that is captured with courser resolution, such as Visium.

Slide-seq is another popular sST technology which uses Drop-seq like beads for oligonucleotide synthesis as spots with diameters of 10µm that are randomly arranged on the array surface (8). A recent updated version of this protocol (Slide-seq V2) modifies the library generation, bead synthesis, and array indexing steps to improve the mRNA capture sensitivity by approximately 10-fold over the original Slideseq method (9). Curio-seeker is the commericially available version of Slide-seq V2.

These three technologies differ significantly in their approaches to spatial barcode synthesis, which directly impacts data quality and the preprocessing strategies. 10x Visium uses chemical or optical methods to directly synthesize and attach barcodes to predefined grid points on the glass slide. Each grid point has a pre-designed unique barcode, which is consistent across different slides, facilitating standardized cross-experiment analysis. BGI Stereo-seq employs Rolling Circle Amplification (RCA) technology to amplify pre-designed barcodes into DNA nanoballs which are densely packed on a solid surface through chemical bonding or electrostatic interactions. Each nanoball carries a unique barcode, distributed with ultra-high density and randomness. This approach supports higher spatial resolution but requires high-precision imaging techniques to decode barcode positions and obtain spatial information. Slide-seq uses chemical coupling reactions to attach pre-designed barcodes onto the surfaces of microbeads, ensuring that each microbead carries a unique barcode. These microbeads are randomly distributed on a substrate, such as a polyacrylamide gel or a glass surface. The spatial positions of the microbeads are resolved through high-resolution imaging techniques, such as fluorescence microscopy, and correlated with their respective barcode sequences. Barcode distribution is thus entirely random.

The systematic differences between sST technologies has led to the development of platform-specific data preprocessing tools, which handle the raw sequencing reads and summarise these into a per sample spatial gene count matrix. For instance, `Space Ranger`, developed by 10x Genomics, processes raw 10X Genomics Visium data alongside bright-field or fluorescence microscopy images (10). The Stereo-seq Analysis Workflow, `SAW`, designed by BGI, facilitates the preprocessing of Stereo-seq data to generate spatial gene expression matrices (11). The Curio Seeker bioinformatics pipeline developed by Curio Bioscience is written in Nextflow to support the preprocessing of sST data from the Curio Seeker spatial mapping kit (12). Notably, this pipeline is compatible with both Singularity (Apptainer) and Docker environments, enabling flexible execution. In terms of open-source analysis solutions for sST data preprocessing, few solutions exist. `Spacemake` is one tool previously developed for use with Visium and Slide-seq data, but it lacks support for nanoball-based ST methods such as BGI Stereo-seq (13). To fill this gap, we developed the `stPipe` R package, which builds upon the foundations laid by our earlier `scPipe` package (14). `stPipe` is the first fully integrated R package that can process paired-end (PE) sequencing reads from multiple sST platforms to create a spatial count matrix in preparation for downstream analysis. The output of `stPipe` is fully compatible with current popular mainstream tools for ST analysis available from various open-source projects, which facilitates a comprehensive beginning to end analysis of sST data. In this article, we introduce the main features and implementation details of `stPipe` and demonstrate its use in quality control and downstream analysis using various bench-marking datasets to compare different technologies.

## Design and implementation

### stPipe implementation details

`stPipe` is an R package that can handle data generated from popular sST protocols, including 10x Visium, BGI Stereo-seq, Slide-seq and Curio-seeker (Figure 1). The pipeline initiates with paired-end (PE) FASTQ files and outputs a gene count matrix with matched spatial locations, quality control (QC) statistics, clustering results, and a summary HTML report. `stPipe` re-uses and extends functionality from a number of other R and python based packages, including `scPipe` for FASTQ reformatting, exon mapping, barcode demultiplexing, and gene counting, `Rsubread` for aligning reads to a reference genome or collection of probe-sets (15), `DropletUtils` to perform QC filtering (16), `Shiny` for the creation of a interactive region of interest (ROI) selection tool (17) and `Seurat` (18), `SpatialExperiment` (19) and `anndata` (20) to organise the spatial counts and sample annotation information. Computationally intensive functions are implemented in C++ and wrapped as R functions using the `Rcpp` package (21). A configuration file is required by `stPipe` to allow users to specify parameters related to the data type being analyzed. For example, paths to FASTA and gff3 files are required for processing samples generated using poly-A-based protocols and paths to h5 mapping files are needed for Stereo-seq based data. The individual function help pages as well as the `stPipe` package vignette provides explanations of various configuration file parameters.

**Fig. 1.**
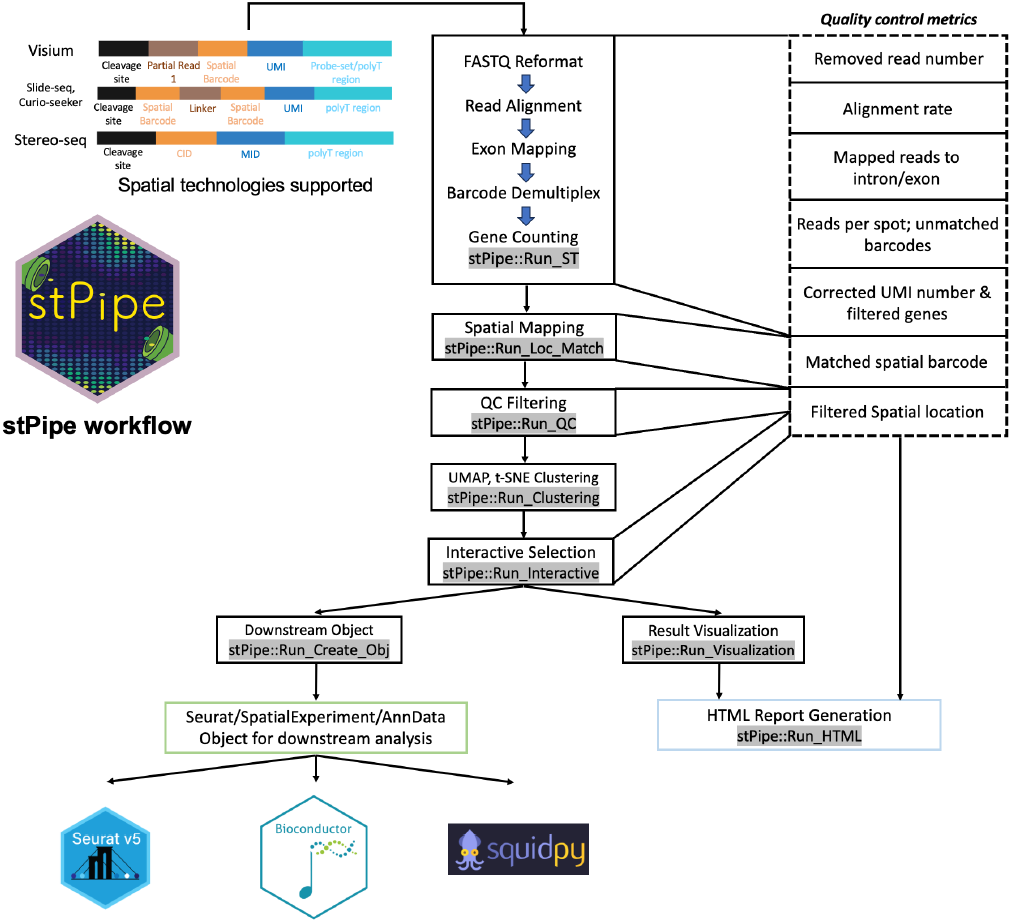
Overview of the stPipe workflow. The major steps in the `stPipe` data preprocessing pipeline are shown along with the QC statistics collected at each stage. The final output of the pipeline is a matrix of gene counts annotated with corresponding spatial information, sample annotations and collected QC metrics during corresponding processing steps for use in downstream analysis and an HTML report that summarizes these data quality metrics. The resulting `Seurat`, `SpatialExperiment` or `Anndata` object can be passed as input to `Seurat` or various Bioconductor packages which rely on `SpatialExperiment` objects, or a `Squidpy` workflow to perform further downstream analysis to explore the biological questions of interest.

### Sequencing-based Spatial Transcriptomics datasets analysed

#### 10x Visium mouse spleen dataset

The *SpatialBenchVisium* dataset (22) provides spatially resolved transcriptomic profiles of mouse spleens from 8-week-old mice post recovery from malaria infection. Samples are available across different combinations of 10x Visium sample handling and profiling protocols, including sample preparation protocols for fresh frozen at optimal cutting temperature (OCT) or formalin-fixed paraffin-embedded (FFPE); tissue placement as direct tissue placement on the slide with the use of CytAssist (CA) or manual placement. Samples can be grouped by sex (male of female) or genotype (knock-out, control or wild-type). The reads were sequenced by an Illumina NextSeq 2000 according to 10x guidelines and processed by `stPipe`. This dataset is available under GEO accession number GSE254652.

#### 10x Visium mouse brain dataset

This dataset originates from the publicly available 10x Genomics Visium platform and represents a coronal section of FFPE mouse brain tissue. Specifically, the sample comprises brain tissue obtained from 8-week-old male mice, oriented in the coronal plane, sectioned to a thickness of 5 µm, and mounted onto Visium Gene Expression slides in accordance with the standardized 10x Genomics Visium protocol. H&E imaging was performed using the Metafer Slide Scanning Microscope (MetaSystems), providing detailed histological context for subsequent spatial gene expression profiling.

#### Slide-seq V2 mouse brain dataset

Mouse hippocampus samples (Puck_191204_01) were analysed using `stPipe` as one of the use cases. A spline was fitted along the CA1 pyramidal cell layer, and beads were averaged to generate a gene expression profile extending approximately 100µm into the basal neuropil and about 400µm into the proximal neuropil (9). Samples were then sequenced based on the Illumina NovaSeq platform. This dataset is available from the Broad Institute’s online single cell portal (23).

#### Stereo-seq mouse brain dataset

This Stereo-seq mouse brain dataset (sample id: SS200000135TL_D1) is publicly available in SRA via BioProject PRJNA1036005 and is used as a demo dataset in the SAW pipeline (11). This sample contains over one billion reads from a chip size of 1 × 1cm.

## Results

### The stPipe workflow

#### Preprocessing of sST data

The `stPipe` workflow (Figure 1) begins with the function `Run_ST` which streamlines multiple steps into one cohesive process, including FASTQ or BAM file reformatting, read alignment, exon mapping, barcode demultiplexing, and gene counting. The `Run_ST` function builds upon the approach followed by `scPipe`, which has been further adapted and extended for the specific requirements of spatial transcriptomics data analysis. Different sST protocols have different configurations of the read structure. Visium stores its 16bp spatial barcode followed by 12bp UMI in FASTQ read 1 while Slide-seq has an 8bp spatial barcode, 18bp bridge sequence, 8bp spatial barcode (7bp in Slide-seq V2) and 8bp UMI (9bp in Slide-seq V2) in FASTQ read 1. Initially, FASTQ or BAM (for Slide-seq based experiments) files are reformatted, where barcodes and UMI information are extracted and incorporated into the read headers. For BGI Stereo-seq (STomics) data, this step involves the mapping and deconvolution of spatial barcodes to their corresponding x,y coordinate pairs as described in the provided mapping file for the chip. This is performed by a C++ script either run standalone or with the provided `Run_ST` function in the `stPipe` package using `Rcpp` (21). Deconvolution is performed on the PE FASTQ files by translating the coordinate and UMI data in read 1 and inserting both into the header for read 2. The resulting FASTQ file is used in the subsequent `stPipe` preprocessing pipeline. Configuration for deconvolution is provided in the required config file including locations of the paired FASTQ files and the coordinate mapping file for the chip.

After reformatting, sequence alignment is performed using `Rsubread`, which maps the reads to a reference genome (or library of probe-sets for Visium probe-based platforms), resulting in a BAM file with positional information. Aligned reads are then assigned to annotated genomic features (e.g. exons or transcripts) using user-provided annotations in GTF or GFF format. Following alignment, exon mapping is performed to assign reads to specific exonic regions. The workflow also includes barcode demultiplexing to identify individual spatial location across the whole tissue section being captured. It is noteworthy that `stPipe` uses different strategies for probe-based and polyA-based protocols. The former uses a probe-set version to construct annotation files such as fa and GFF3 files while the latter relies on the gene annotation files specified by the user.

Lastly, the `Run_ST` function generates a gene count matrix after performing UMI deduplication to remove PCR duplicates. The deduplication approach in `stPipe` uses a distance-based method, comparing UMI sequences within a specified Hamming distance. When one UMI has significantly more reads than the other, the two are considered duplicates. This ensures accurate gene quantification by minimizing the impact of sequencing and PCR errors. Additionally, the `Run_ST` function implements parallel computing via the `mclapply` function in order to save computing time when processing batches of samples.

#### Matching spatial with gene count information

After obtaining the gene count matrix with relevant sample index which refers to the spatial barcode sequence using the `Run_ST` function, the `Run_Loc_Match` function can be used to obtain spatial locations with corresponding UMI counts for different technologies. For 10x Visium, there are currently five different chip types, each of which has a fixed relationship between spatial barcodes and spatial coordinates. For Stereo-seq, Slide-seq, and Curio-seeker, barcodes are randomly distributed across spatial coordinates which must be mapped in a sample-specific way. For 10x Visium data, `stPipe` requires the user to specify the chip version and then uses internally stored 10x Visium coordinates to match the spatial barcodes. `stPipe` also provides additional pixel computation of each spot for Visium technology which is achieved by circle detection based on Python `OpenCV` library (24). For the remaining protocols, users are required to provide a path to the sample-specific spatial coordinate csv file.

#### Filtering out low quality spatial locations

After matching the gene count profile with corresponding spatial locations, the `Run_QC` function can be used to filter out low quality data (Figure 2). `stPipe` provides two different options for this: `max_slope` and `EmptyDropletUtils`. For the former approach, the filtering process is carried out based on the raw UMI count in every spatial location. Locations with counts that fall below a certain threshold are considered low quality and removed from the gene count matrix. This method retains only the spatial locations with significant transcriptomic signals to reduce noise from locations with minimal or no meaningful biological information. The threshold in this method is computed by analyzing the distribution of UMI counts across spatial locations, and identifying the point of maximum slope in the cumulative UMI distribution curve. This point often corresponds to the transition between background noise and biological signal. The second `EmptyDropletUtils` option makes use of the `DropletUtils` package which offers a more sophisticated computation to identify spatial locations that contain real cells, as opposed to positions that contain ambient RNA or other noise. This method calculates a false discovery rate (FDR) to assess the likelihood of each spatial location containing one or more real cells. Filtering can be fine-tuned using both *p*-value and FDR thresholds, offering greater flexibility in distinguishing between noise and meaningful data. Spatial locations are retained if they meet the significance criteria for either the *p*-value or FDR. These two thresholds are specified in the `stPipe` config file.

**Fig. 2.**
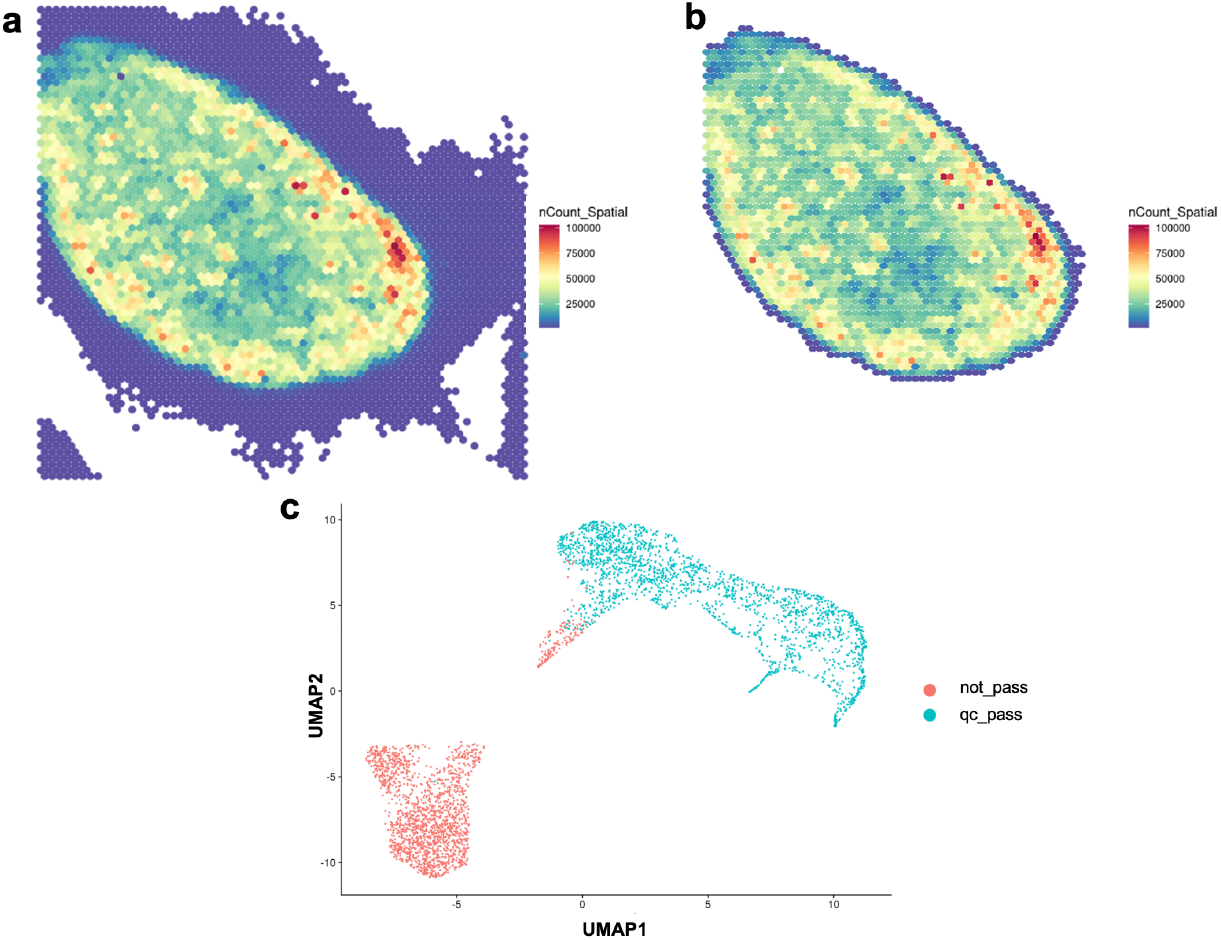
Results before and after the Run_QC function in spatial and cluster views. (a) Spatial heatmap of raw UMI count before the `Run_QC` function of FFPE, CytAssist mouse spleen sample 709. (b) Spatial heatmap of raw UMI count after the `Run_QC` function is applied to this sample. The spatial heatmap suggests the tissue integrity of mouse spleen as well as successful removal of background using the `Run_QC` function. (c) UMAP representation of the FFPE, CytAssist, mouse spleen sample 709 after the `Run_QC` function is applied, spots pass QC filter and spots do not are projected as two colors. This suggests the removal of background from tissue.

#### Interactive selection of regions of interest

The `Run_QC` function primarily acts as a tool for filtering out low-quality data, however it may not effectively eliminate all background or out-of-tissue spots for the Visium platform with high precision. To address this challenge, the `Run_Interactive` function was implemented to allow users to more accurately select regions of interest (ROI) such as in-tissue spots for Visium data. This function is built based on the R `shiny` package (17) with embedded `Plotly` for enhanced interactivity, supporting various selection tools including zoom, pan, rectangular selection, and lasso selection. It also enables linking of each spot’s spatial location to its corresponding point in the interactive UMAP (Uniform Manifold Approximation and Projection) or T-SNE (t-Distributed Stochastic Neighbor Embedding) plots provided by the `Run_Clustering` function. Prior to dimensionality reduction and clustering, highly variable genes (HVGs) are selected. After feature selection, the `Run_Clustering` function performs dimensionality reduction via UMAP and t-SNE, followed by clustering using methods such as Louvain, Leiden, or *k*-means. It thereby produces interactive plots wherein each spot’s spatial location is linked to its corresponding point in the UMAP or t-SNE embedding. This linkage facilitates an intuitive comparison between the reduced dimensional representation, the clustering outcomes, and the original spatial coordinates (Figure 3).

**Fig. 3.**
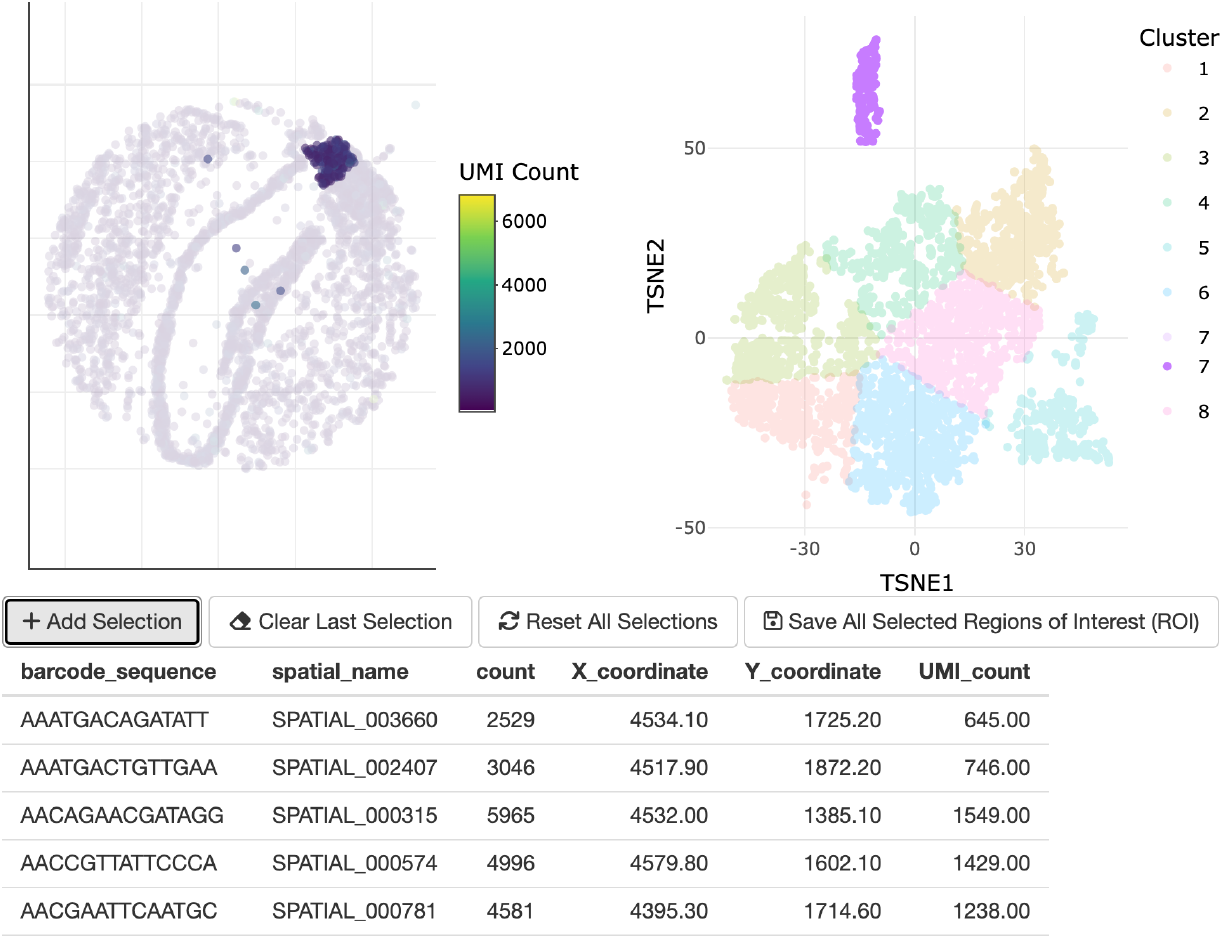
Screenshot of Interactive R-shiny web app created by the Run_Interactive function for Slide-seq mouse brain sample Puck_200115_08. This interactive function offers flexible options for selecting a regions of interest (ROIs) through four intuitive buttons. The “Add Selection” button allows users to add spatial coordinates along with corresponding metadata such as UMI count and spatial barcode sequences each time an ROI is selected. The “Clear Last Selection” button removes the most recently selected ROI from the current selection list. The “Reset All Selections” button resets both the spatial heatmap and clustering plot, providing a clean slate for a new selection. Finally, the “Save All Selected ROI” button saves the finalized selection as “selected_ROI” object in the user’s R global environment, streamlining data management and export. In this example, cluster 7 highlighted in purple on the T-SNE plot has been selected is found to mostly correspond to the choroid plexus region in the spatial UMI count plot.

Users can either select the ROI directly on the spatial image, which is then mapped onto the UMAP or T-SNE plot, or select a specific cluster on the UMAP or T-SNE clustering map, with the selected cluster points mapped back to their corresponding spatial locations. In addition, the `Run_Interactive` function includes four interactive buttons which include “Add Selection”, “Clear Last Selection”, “Reset All Selections”, and “Save All Selected ROI”, allowing users to easily manage their selections. As the selected ROIs are visually highlighted, users can precisely choose regions of interest and easily save the refined data back to the global environment in R, enhancing control and accuracy in data selection.

### Data visualization and quality assessment

The `Run_Visualization` function visualizes both spatial-level and read-level information for the results generated in the previous steps of `stPipe`. This function outputs both raw and log-transformed UMI counts for spatial visualization in a heatmap format (Figure 2a-b), while demultiplexing results are shown in barplot format (Figure 4a), mapping statistics as a stacked barplot (Figure 4b) and UMI duplication number as a line plot (Figure 4c). For multi-sample experiments, UMI counts distribution plots can be produced to assess data quality between samples (Figure 4d). These metrics provide complementary insights into both spatial-level and read-level quality. For example, lower exon mapping rates or high ambiguity in alignment can indicate suboptimal RNA quality or technical issues during library preparation, while excessive PCR duplication revealed by UMI deduplication statistics may indicate low library complexity. The spatial UMI heatmap can flag tissue integrity issues or specific spatial patterns when compared to the corresponding H&E image. Overall, these QC metrics are valuable for flagging poor-quality samples or tissue regions to filter out before proceeding to downstream analysis.

**Fig. 4.**
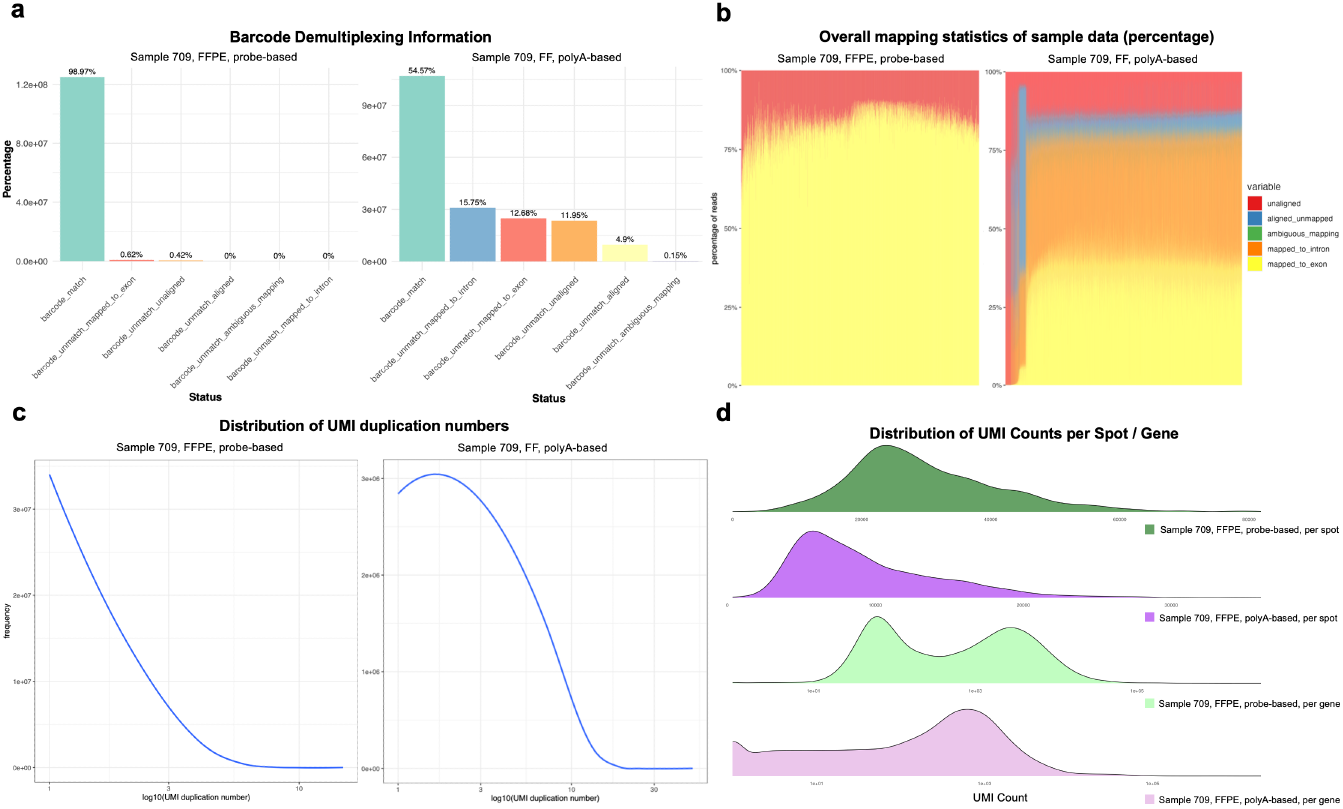
Visualization results generated by stPipe for data quality assessment. (a) Barplot showing spatial barcode demultiplexing information between 10x Visium probe-based (left) and polyA-based (right) protocols to assess sequencing accuracy. (b) Stacked bar plots showing the mapping rate, separated into reads that map to exons, introns, and those that are ambiguously mapped, map elsewhere in the genome (ordered by exon mapping rate) between 10x Visium probe-based (left) and polyA-based (right) protocols. (c) UMI duplication plot between a probe-based sample (left) with a higher UMI duplication number than a polyA-based one (right). A distribution skewed toward lower duplication values indicates higher library complexity and minimal redundancy, suggesting that the sequencing depth is well-matched to the diversity of the transcriptome. In contrast, a pronounced tail toward higher duplication values suggests substantial over-sequencing or PCR amplification biases, as many reads may originate from the same underlying transcript molecule. (d) UMI count distribution between sample 709 with two protocols, the first and last two are plotted as distribution of raw UMI count per spot and log_10_ UMI count per gene respectively. Different protocols are shown in distinct colours.

### Automatic generation of HTML report

The `Run_HTML` function is designed to automatically generate an HTML report in R Markdown format. This function takes outputs from the `Run_QC`, `Run_Visualization` and `Run_Clustering` functions and builds a report with plots summarising barcode demultiplexing, UMI duplication, mapping statistics, QC thresholds, the number of spatial locations before and after the `Run_QC` step, and interactive T-SNE and UMAP plots with clustering results, each within its own subsection to illustrate its relevant importance within the pipeline. A key feature of `stPipe` is the collection of a consistent set of QC measures and data visualizations across different platforms which facilitate comparisons between multiple datasets.

### Object creation for downstream analysis

Spatial count data is organised by the `Run_Create_Obj` function which constructs flexible objects for downstream analysis across various mainstream workflows. `Run_Create_Obj` allows users to create: (1) R `Seurat` spatial objects to allow compatibility with `Seurat` sST workflows; (2) R `SpatialExperiment` objects for use in Bioconductor sST workflows such as `nnSVG` for Spatially Variable Gene (SVG) identification (25) and `STdeconvolve` (26) or `SpatialDecon` (27) for cell type inference and deconvolution; and (3) Python `AnnData` spatial objects for use in `Squidpy` sST workflows (20).

### Config file

`stPipe` simplifies the use of each function and overall code readability via a config file. The essential inputs in the config file include the data directory, output directory, sample species, technology version, specified spatial coordinate system, the format of the read structure, number of threads to process data over, number of reads to process and the number of spatial locations. Other platform-specific information such as the spatial location csv file path for Slideseq or Curio-seeker data, specific fa and gff files for polyA-based sequencing methods, tiff image path for pixel computation, path to h5 mapping file for Stereo-seq data and the QC method and the ratio set for `Run_QC` function. When dealing with multiple samples, only those run using the same platform and species can be processed together in `stPipe`.

## stPipe use cases

A vignette accompanying the package provides further details on implementation and an example use case on the downsampled 10x Visium FFPE, CytAssist processed mouse spleen sample 709 from the `SpatialBenchVisium` dataset (22). Several other use cases are presented in the following sections.

### Analysis of a 10x Visium mouse spleen dataset

Multiple samples (167, 168, 544, 545, 708 and 709) from the *SpatialBenchVisium* dataset (22) were preprocessed using `stPipe`. Each sample contains over 10 million reads and takes under 30 minutes to process on a standard Linux server using the current `stPipe` workflow for either the polyA-based or probe-based Visium platform. After spatial location matching and selection, the spots retained for each sample by `stPipe` shared over 98% intersection ratio with the 10x `Space Ranger` output. For each sample, UMI count and captured gene per spot had very similar distributions between `stPipe` and 10x `Space Ranger` outputs, indicating `stPipe` recovers comparable results to the vendor provided software for Visium data preprocessing.

The `Run_Create_Obj` function, followed by downstream analyses that included spatially variable gene (SVG) detection, differential expression (DE) analysis, ROAST gene set testing (28), and cell type inference, followed the same workflow outlined in the original study by Du *et al*. (22). The downstream results from gene-specific (SVG, DE) and cell-type-specific analyses further underscore the reliability of `stPipe`. Specifically, the *MA*-plot for the DE comparison of the B cell cluster (see Supplementary Figure S2C) reveals a comparable trend in DE analysis to that reported in the original study which was based on `Space Ranger` output, with the accurate detection of enrichment of sex-specific genes (those from the Y chromosome and genes on the X chromosome that escape X inactivation) that are expected to differ when comparing male versus female wild-type samples (ROAST *p*-value = 0.0005). These results affirm `stPipe`’s capability to consistently identify key biological features. In terms of SVG and HVG identification, *Car2* consistently emerged as the top-ranked gene in both categories, aligning with the findings of the original paper. Additionally, cell type inference results (Supplementary Figure S3) show comparable proportions, with an average difference of less than 2% across cell types when comparing `stPipe` with `Space Ranger` results. Both analyses employed an identical marker gene list, covering plasma cells, T cells, neutrophils, B cells, germinal centers and erythrocytes, which underscores the robustness and consistency in cell type classification between these two preprocessing pipelines.

### Analysis of mouse brain datasets across Visium, Slide-seq, and Stereo-seq platforms

Publicly available datasets from Visium, Slide-seq, and Stereo-seq were preprocessed from raw FASTQ or BAM files into gene count matrices using `stPipe` with corresponding workflows implemented in the `Run_ST` function. Preprocessing these data on a standard Linux server took around 30 minutes for 65 million PE reads from the Visium sample, 2 hours for over 200 million PE reads from the Slide-seq sample and 8 hours for the 270 million PE reads from the Stereo-seq sample. Next, the `Run_Loc_Match` function was used to obtain matched spatial location information. Finally, `Seurat` spatial objects were constructed via the `Run_Create_Obj` function, followed by downstream analyses which includes marker gene identification, and inference of both cell types and spatial domains using the `Seurat` workflow.

In the mouse brain, the hippocampal region CA2 is distinguished from the CA1 and CA3 regions by several unique characteristics, such as its distinct gene expression profile, lack of long-term potentiation, and higher resistance to cell death. However, due to the hippocampus’s curved structure, accurately locating CA2 relative to CA1 and CA3 along the transverse axis within coronal and sagittal slices, as shown in the Allen Brain Atlas (29), can be challenging. Hence, the result of spatial domain identification of the CA2 region with related significant marker genes can be considered as an important indicator of `stPipe`’s preprocessing performance. In addition to inferring cell types and spatial domains, marker gene sensitivity tests have been performed to validate these inference outcomes via the `FindMarkers` function implemented in the `Seurat` package with a significance criteria of log-fold change greater than 0.25 and adjusted *p*-value less than 0.01. Domain-specific marker genes includes *Hpca, Prox1, Ptgds, Ttr, Prdm8*, and *Slc17a7* for the hippocampus region and region-specific marker genes including *Rgs14, Camk4, Amigo2*, and *Ntf3* for the CA2 region were used for evaluation.

Among these platforms, Visium (Figure 5a) offers lower spatial resolution but captures broad regional differences, making it suitable for identifying general spatial patterns; Slideseq (Figure 5b) improves upon resolution, enabling the identification of finer hippocampus structures with the CA2 region inside it; Stereo-seq (Figure 5c) provides the highest resolution among the three platforms, effectively distinguishing sub-regional domains, such as the CA1 and CA3 regions, but it fails to identify the CA2 region accurately. For marker gene sensitivity testing, most of the significant domain-specific and region-specific marker genes were consistently captured by three platforms for most regions, except for Visium and Stereo-seq which failed to detect CA2 marker genes across the captured spatial section.

**Fig. 5.**
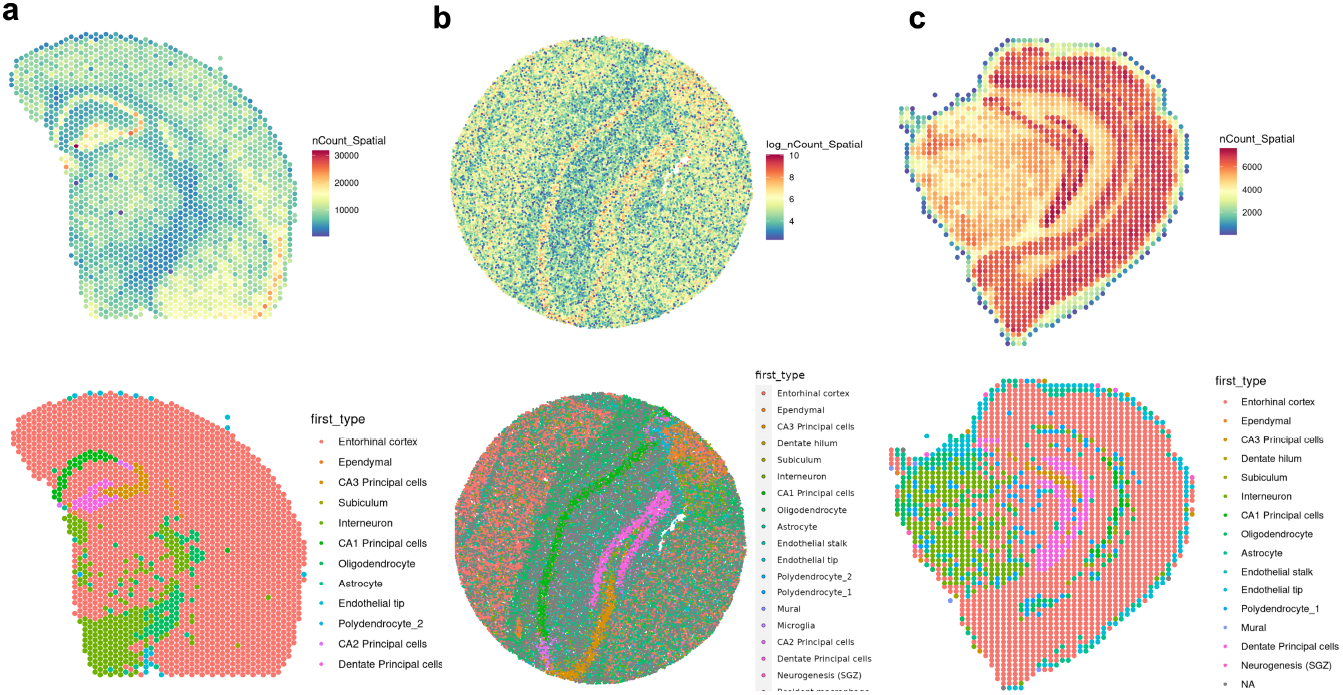
Spatial UMI plot and deconvolution result of mouse brain across different platforms. (a) Visualization of mouse brain sample based on the 10x Visium platform. The upper panel depicts the spatial UMI distribution following preprocessing with the `stPipe` workflow, and the lower panel illustrates the deconvolution results obtained using the RCTD algorithm implemented in the `Seurat` workflow; (b) Visualization of mouse brain sample processed based on Slide-seq platform, following the same figure structure and workflow as in (a); (c) Visualization of mouse brain sample processed based on Stereo-seq platform, following the same figure structure and workflow as in (a); Further details regarding the datasets and analyses are provided in the dataset description and analysis sections, respectively.

## Availability and future directions

The `stPipe` package is available from https://github.com/mritchielab/stPipe and has been submitted to Bioconductor. It has passed the pre-check status and undergoing review process. Code for the examples described in the ‘**stPipe use cases**’ section above is available in the package vignette and the remaining analysis code for this paper is available from GitHub at https://github.com/YangXuuu/stPipe_manuscript.

Despite the rapid growth of the ST field across a diverse range of technology platforms and data analysis tools tailored to ST analysis, there are limited options for handling the raw data obtained from mainstream sST protocols from beginning to end and collecting detailed QC information in a unified way. The `stPipe` package bridges this gap, accepting raw FASTQ files and allowing flexiblity in read structure to support data output by various sST protocols, including 10x Visium, Slide-seq and Stereo-seq. `stPipe` outputs numerous QC metrics obtained during the data preprocessing workflow and presents these results in an HTML report to assist users in QC evaluation and downstream analysis. `stPipe` is an being further developed, with planned future improvements in the following three areas: 1. including support for various downstream analysis tasks such as SVG identification, deconvolution/cell type inference, marker gene detection, and cell-cell communication based on the results obtained from upstream outputs; 2. application of `stPipe` in larger scale sST benchmarking analyses; 3. extending support to recently released sST platforms such as 10x Visium HD and BGI Stereo-seq with grouped FASTQ file format.

## Acknowledgments

The authors are grateful to Kathleen Zeglinski for creating the `stPipe` package logo.

## Funding

This work was supported by funding from the Australian National Health and Medical Research Council (NHMRC) Investigator Grant 2017257 to M.E.R.

## Competing interests

The authors declare that they have no competing interests.

## Authors’ contributions

Y.X. developed software, performed data analysis, generated figures and wrote the manuscript. C.J.S., Y.Y., Y.Y, S.S. and C.W. contributed to software development and data analysis. L.T., Y.C, and M.E.R. designed the study, supervised data generation, analysis and interpretation and wrote the manuscript. All authors read and approved the final manuscript.

## Abbreviations

BP: Base pair;
CID: Coordinate Identity;
DEG: Differentially Expressed Gene;
DNB: DNA Nanoball;
FDR: False Discovery Rate:
FF: Fresh Frozen;
FFPE: Formalin Fixed Paraffin-Embedded;
HE: Hematoxylin and Eosin;
HVG: Highly Variable Gene;
iST: Image-based Spatial Transcriptomics;
NGS: Next Generation Sequencing;
PE: Paired end;
QC: Quality control;
ROI: Region of interest;
RTL: RNA templated ligation;
SAW: Stereo-seq Analysis Workflow;
scRNA-seq: Single-Cell RNA Sequencing;
sST: Sequencing-based Spatial Transcriptomics;
Stereo-seq: SpaTial Enhanced REsolution Omics-sequencing;
ST: Spatial Transcriptomics;
SVG: Spatially Variable Gene;
T-SNE: t-Distributed Stochastic Neighbor Embedding;
UMAP: Uniform Manifold Approximation and Projection;
UMI: Unique Molecular Identifier

## Notes

https://singlecell.broadinstitute.org/single_cell/study/SCP815/highly-sensitive-spatial-transcriptomics-at-near-cellular-resolution-with-slide-seqv2

https://github.com/mritchielab/stPipe

https://www.ncbi.nlm.nih.gov/geo/query/acc.cgi?acc=GSE254652

### Competing Interest Statement

The authors have declared no competing interest.

## References

1. Vivien Marx. Method of the year: spatially resolved transcriptomics. Nature methods, 18(1):9–14, 2021.

2. Ye Wang, Bin Liu, Gexin Zhao, YooJin Lee, Anton Buzdin, Xiaofeng Mu, Joseph Zhao, Hong Chen, and Xinmin Li. Spatial transcriptomics: Technologies, applications and experimental considerations. Genomics, page 110671, 2023.

3. Lambda Moses and Lior Pachter. Museum of spatial transcriptomics. Nature methods, 19(5):534–546, 2022.

4. Luyi Tian, Fei Chen, and Evan Z Macosko. The expanding vistas of spatial transcriptomics. Nature Biotechnology, 41(6):773–782, 2023.

5. Patrik L Ståhl, Fredrik Salmén, Sanja Vickovic, Anna Lundmark, José Fernández Navarro, Jens Magnusson, Stefania Giacomello, Michaela Asp, Jakub O Westholm, Mikael Huss, et al. Visualization and analysis of gene expression in tissue sections by spatial transcriptomics. Science, 353(6294):78–82, 2016.

6. Michelli F Oliveira, Juan P Romero, Meii Chung, Stephen Williams, Andrew D Gottscho, Anushka Gupta, Susan E Pilipauskas, Syrus Mohabbat, Nandhini Raman, David Sukovich, et al. Characterization of immune cell populations in the tumor microenvironment of colorectal cancer using high definition spatial profiling. bioRxiv, pages 2024–06, 2024.

7. Ao Chen, Sha Liao, Mengnan Cheng, Kailong Ma, Liang Wu, Yiwei Lai, Xiaojie Qiu, Jin Yang, Jiangshan Xu, Shijie Hao, et al. Spatiotemporal transcriptomic atlas of mouse organogenesis using dna nanoball-patterned arrays. Cell, 185(10):1777–1792, 2022.

8. Samuel G Rodriques, Robert R Stickels, Aleksandrina Goeva, Carly A Martin, Evan Murray, Charles R Vanderburg, Joshua Welch, Linlin M Chen, Fei Chen, and Evan Z Macosko. Slide-seq: A scalable technology for measuring genome-wide expression at high spatial resolution. Science, 363(6434):1463–1467, 2019.

9. Robert R Stickels, Evan Murray, Pawan Kumar, Jilong Li, Jamie L Marshall, Daniela J Di Bella, Paola Arlotta, Evan Z Macosko, and Fei Chen. Highly sensitive spatial transcriptomics at near-cellular resolution with slide-seqv2. Nature biotechnology, 39(3):313–319, 2021.

10. Amanda Janesick, Robert Shelansky, Andrew D Gottscho, Florian Wagner, Stephen R Williams, Morgane Rouault, Ghezal Beliakoff, Carolyn A Morrison, Michelli F Oliveira, Jordan T Sicherman, et al. High resolution mapping of the tumor microenvironment using integrated single-cell, spatial and in situ analysis. Nature Communications, 14(1):8353, 2023.

11. Chun Gong, Shengkang Li, Leying Wang, Fuxiang Zhao, Shuangsang Fang, Dong Yuan, Zijian Zhao, Qiqi He, Mei Li, Weiqing Liu, et al. Saw: An efficient and accurate data analysis workflow for stereo-seq spatial transcriptomics. GigaByte, 2024, 2024.

12. Curio seeker spatial transcriptomics kits. https://curiobioscience.com/seeker/. 2024.10.20.

13. Tamas Ryszard Sztanka-Toth, Marvin Jens, Nikos Karaiskos, and Nikolaus Rajewsky. Spacemake: processing and analysis of large-scale spatial transcriptomics data. Gigascience, 11:giac064, 2022.

14. Luyi Tian, Shian Su, Xueyi Dong, Daniela Amann-Zalcenstein, Christine Biben, Azadeh Seidi, Douglas J Hilton, Shalin H Naik, and Matthew E Ritchie. scpipe: A flexible r/bioconductor preprocessing pipeline for single-cell rna-sequencing data. PLoS computational biology, 14(8):e1006361, 2018.

15. Yang Liao, Gordon K Smyth, and Wei Shi. The subread aligner: fast, accurate and scalable read mapping by seed-and-vote. Nucleic acids research, 41(10):e108–e108, 2013.

16. Aaron TL Lun, Samantha Riesenfeld, Tallulah Andrews, The Phuong Dao, Tomas Gomes, Participants in the 1st Human Cell Atlas Jamboree, and John C Marioni. Emptydrops: distinguishing cells from empty droplets in droplet-based single-cell rna sequencing data. Genome biology, 20:1–9, 2019.

17. Winston Chang, Joe Cheng, J Allaire, Yihui Xie, Jonathan McPherson, et al. Shiny: web application framework for r. R package version, 1(5):2017, 2017.

18. Yuhan Hao, Tim Stuart, Madeline H Kowalski, Saket Choudhary, Paul Hoffman, Austin Hartman, Avi Srivastava, Gesmira Molla, Shaista Madad, Carlos Fernandez-Granda, et al. Dictionary learning for integrative, multimodal and scalable single-cell analysis. Nature biotechnology, 42(2):293–304, 2024.

19. Dario Righelli, Lukas M Weber, Helena L Crowell, Brenda Pardo, Leonardo Collado-Torres, Shila Ghazanfar, Aaron TL Lun, Stephanie C Hicks, and Davide Risso. Spatialexperiment: infrastructure for spatially-resolved transcriptomics data in r using bioconductor. Bioinformatics, 38(11):3128–3131, 2022.

20. Giovanni Palla, Hannah Spitzer, Michal Klein, David Fischer, Anna Christina Schaar, Louis Benedikt Kuemmerle, Sergei Rybakov, Ignacio L Ibarra, Olle Holmberg, Isaac Virshup, et al. Squidpy: a scalable framework for spatial omics analysis. Nature methods, 19(2):171–178, 2022.

21. Dirk Eddelbuettel and Romain François. Rcpp: Seamless r and c++ integration. Journal of statistical software, 40:1–18, 2011.

22. Mei RM Du, Changqing Wang, Charity W Law, Daniela Amann-Zalcenstein, Casey JA Anttila, Ling Ling, Peter F Hickey, Callum J Sargeant, Yunshun Chen, Lisa J Ioannidis, et al. Spotlight on 10x visium: a multi-sample protocol comparison of spatial technologies. bioRxiv, pages 2024–03, 2024.

23. Murray E. Kumar P. Stickels, R.R. et al. Highly sensitive spatial transcriptomics at near-cellular resolution with slide-seqv2. https://singlecell.broadinstitute.org/single_cell/study/SCP815/sensitive-spatial-genome-wide-expression-profiling-at-cellular-resolution#study-summary. 2024.05.20.

24. G. Bradski. The OpenCV Library. Dr. Dobb’s Journal of Software Tools, 2000.

25. Lukas M Weber, Arkajyoti Saha, Abhirup Datta, Kasper D Hansen, and Stephanie C Hicks. nnsvg for the scalable identification of spatially variable genes using nearest-neighbor gaussian processes. Nature communications, 14(1):4059, 2023.

26. Brendan F Miller, Feiyang Huang, Lyla Atta, Arpan Sahoo, and Jean Fan. Reference-free cell type deconvolution of multi-cellular pixel-resolution spatially resolved transcriptomics data. Nature communications, 13(1):2339, 2022.

27. Patrick Danaher, Youngmi Kim, Brenn Nelson, Maddy Griswold, Zhi Yang, Erin Piazza, and Joseph M Beechem. Advances in mixed cell deconvolution enable quantification of cell types in spatial transcriptomic data. Nature communications, 13(1):385, 2022.

28. Di Wu, Elgene Lim, François Vaillant, Marie-Liesse Asselin-Labat, Jane E Visvader, and Gordon K Smyth. Roast: rotation gene set tests for complex microarray experiments. Bioinformatics, 26(17):2176–2182, 2010.

29. Quanxin Wang, Song-Lin Ding, Yang Li, Josh Royall, David Feng, Phil Lesnar, Nile Graddis, Maitham Naeemi, Benjamin Facer, Anh Ho, et al. The allen mouse brain common coordinate framework: a 3d reference atlas. Cell, 181(4):936–953, 2020.

