## Supplementary Table and Figures for "*stPipe:* A flexible and streamlined R/Bioconductor pipeline for preprocessing sequencing-based spatial transcriptomics data"

| **Cell type/splenic region** | **Selected marker genes** |
|---|---|
| B cell | *Cd19, Cd22, Ighd, Cd5* |
| T cell | *Trac, Cd3d, Cd4, Cd3e, Cd8a* |
| Macrophage | *Cd274, Marco, Csf1r, Adgre1, Cd209b, Cd206, Cd80, Mac1, Cd68* |
| Neutrophil | *S100a9, S100a8, Ngp* |
| Erythrocyte | *Car2, Car1, Klf1* |
| Plasma cell | *Cd38, Cd138, Xbp1, Irf4, Prdm1, Cd27, Cd319, Mum1* |
| Germinal centre | *Cxcr4, Cd83, Bcl6, Rgs13, Aicda* |
| Marginal zone | *Marco, Lyz2, Ighd, Igfbp7, Igfbp3, Ly6d* |

**Table S1.** A list of marker genes for cell types or different tissue regions expected in the mouse spleen selected from previous studies and existing literature. These marker genes were used to compute a "cluster score," calculated as the $\log_2$fold-change in expression between each spatial cluster and all other clusters. This quantitative measure was then used to inform the cell type annotation for spatial clusters.

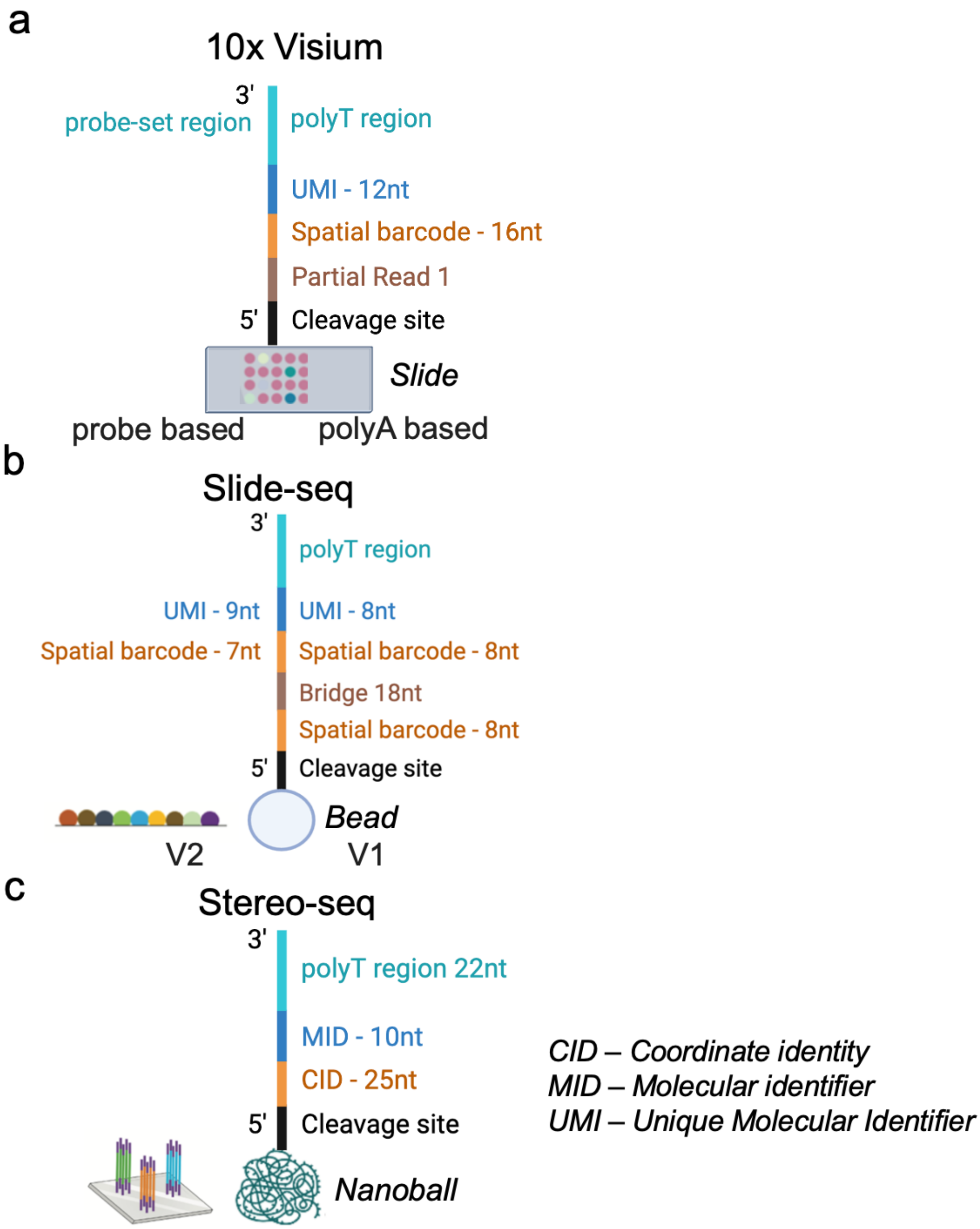

**Figure S1.** Figure presenting FASTQ sequence configuration details across different technologies, including 10x Visium, Slide-seq, and BGI Stereo-seq.

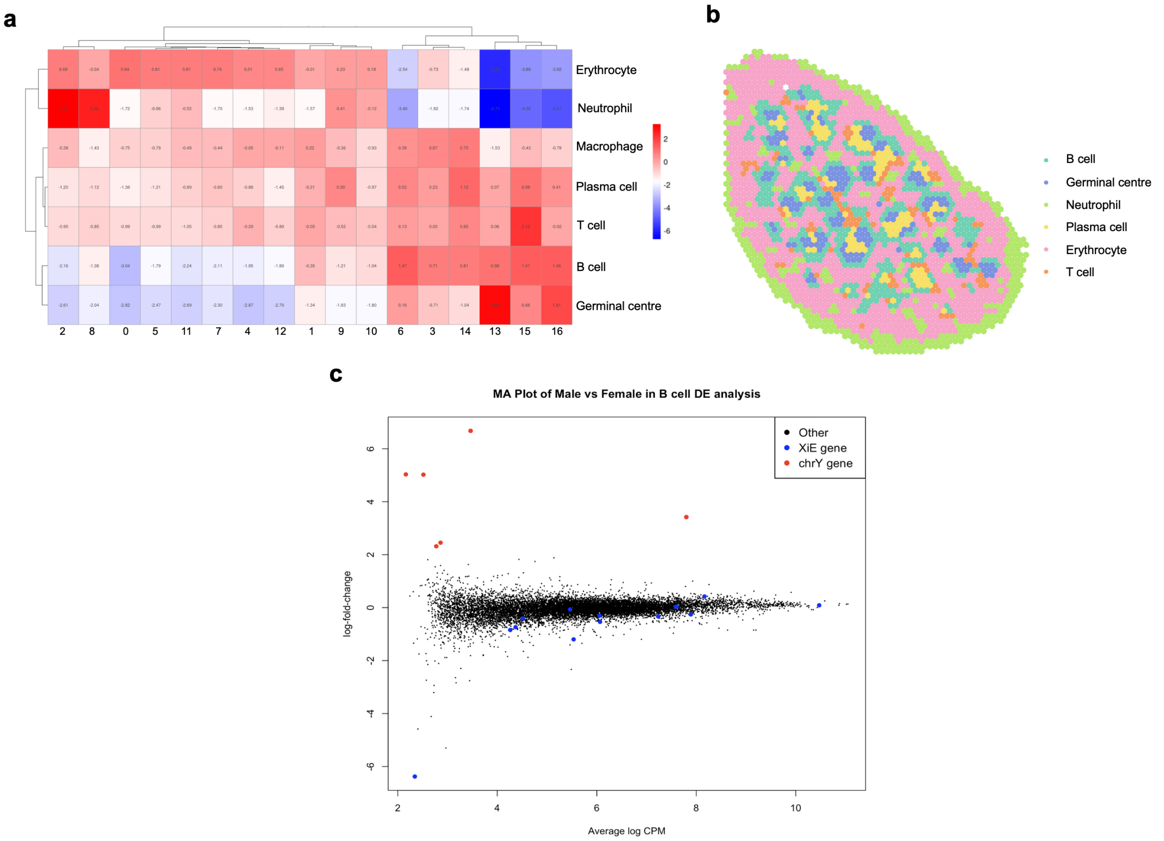

**Figure S2.** Plot showing the intermediate results of analysing the 10x Visium mouse spleen data. (a) Heatmap of cluster vs. cell type with corresponding log-2 fold-change; (b) Spatial dim plot of mouse spleen, showing spatial distribution of B cell, Germinal centre, Neutrophil, Plasma cell, Red pulp, and T cell. (c) $MA$-plot showing sex-specific genes expected to be different between male and female samples in the B cell DE analysis, such as genes on the X chromosome that escape X inactivation (blue points) and chromosome Y genes (red points).

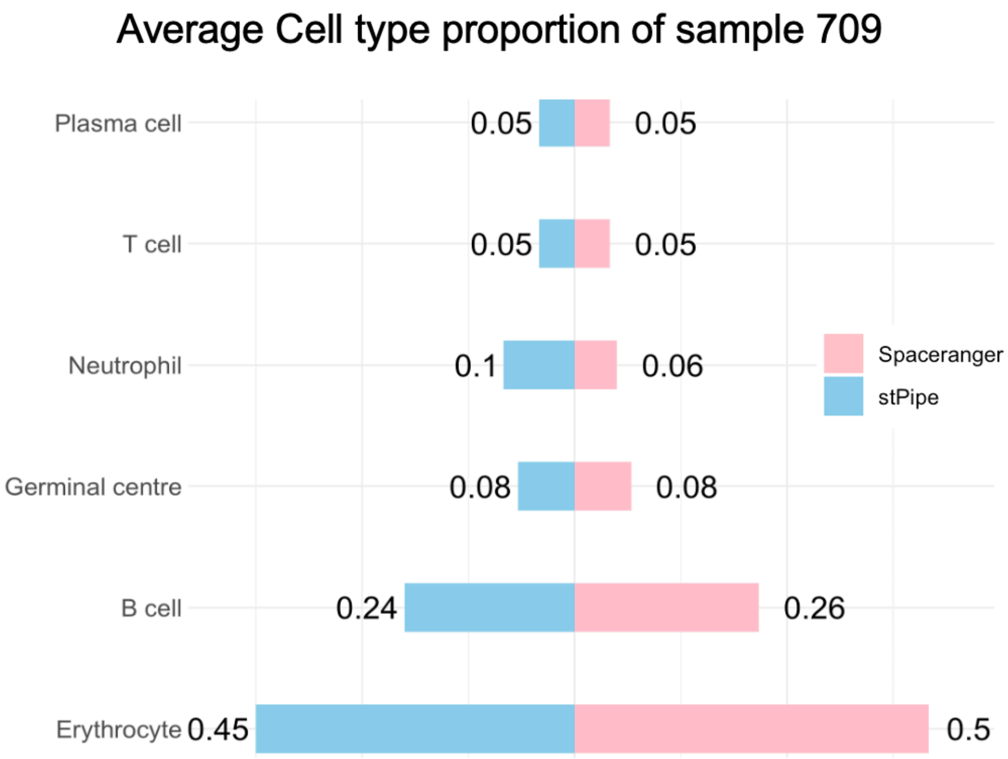

**Figure S3.** Plot showing the average cell type proportion of mouse spleen sample 709 across 4 Visium protocols, grouped by cell type in the y-axis for the different preprocessing tools `Space Ranger` (pink) and `stPipe` (light blue).
